# *Curcuma longa* Essential Oil Topically Mitigates Inflammatory Markers in Acetone-Induced Atopic Dermatitis in Wistar Albino Rats

**DOI:** 10.1101/2024.12.14.628498

**Authors:** Nathiim Namale, Deusdedit Tusubira, Kenneth Male, Angela Mumbua Musyoka, Patrick Maduabuchi Aja

## Abstract

**Background:** Atopic dermatitis ranks number one in skin diseases worldwide. It is challenging to treat; associated with burdens such as sleeping difficulties, depression, and anxiety and impairs life quality due to finances. This study aims to evaluate the potential of topical *Curcuma Longa* Essential Oil in the management of atopic dermatitis. It was driven by overwhelming side effects that present with the use of topical corticosteroids in the management of the condition yet they present a variety of side effects.

**Methods:** *Curcuma Longa* Essential Oil was extracted by hydro-distillation of fresh rhizomes in a Clevenger apparatus. Thirty Wistar Albino rats; Male and Female; were divided into 5 groups, n=6. Atopic dermatitis was induced by rubbing a cotton ball soaked in acetone on rat skin for 5 minutes for three days. Groups were treated with *Curcuma Longa E*ssential oil (1), 0.1% hydrocortisone (2), both *Curcuma Longa* Essential oil and hydrocortisone (3) and the remaining two groups were positive and negative control. The effect of *Curcuma Longa* Topical Essential oil was analysed using its effect on pruritus, Mast cell infiltration in skin biopsies and CRP levels in serum.

**Results:** Pruritus decreased with the *Curcuma Longa* essential oil (*CL*EO) topical treatment and significantly compared to that in the positive group (P <0.0001.). Negative Control Vs *Curcuma longa* essential oil treatment showed a significant difference (P = 0.0001) in CRP levels. Mast cells significantly decreased in groups that had treatment compared to the Positive control.

**Conclusion:** *Curcuma longa* Topical essential oil has the potential treat Acetone induced Atopic Dermatitis in Wistar Albino rats.

**Recommendation:** 2% *Curcuma Longa* topical Essential oil on Atopic dermatitis should be studied on a Clinical trial.

## Introduction

Atopic dermatitis (AD) is number one skin disease worldwide affecting 2.6% population (Tian et al., 2024) with 13.5% in Uganda (Mpairwe et al., 2021). It is an interaction between skin barrier impairment and an immune response with inflammation characterized by a thin epidermis, poor hydration, raised trans-epidermal water loss (TEWL) and increased permeability (Douladiris et al., 2023). It presents with lesions in the acute phase and lichenification in the chronic phase (Al-Afif et al., 2019). Clinically, the hallmark characteristic is a dry itchy skin (Wollenberg et al., 2023). Its incidence has risen in many developing countries due to changes in life style, nutrition and environmental factors (Civelek et al., 2011). Despite the high prevalence of AD in the Uganda population, its management is largely based on the application of Topical corticosteroids hydrocortisone and betamethasone which are characterised by long-term side effects (Mastan, 2020) thus the need for alternative remedies with less and/or more manageable side effects.

*Curcuma longa* (*CL*) has been used as a medicinal herb for the treatment of inflammation since ancient years (Singletary, 2020). It treats psoriasis (Umapathy et al., 2022) an inflammatory skin disease and has therapeutic potential on skin pruritus (Mata et al., 2021). This makes it a good candidate for managing AD an inflammatory skin disease with pruritus. The therapeutic benefits of *CL* are attributed to bioactive compounds curcumin and aromatic turmerone, present in its rhizome (Aydin et al., 2024). The therapeutic benefits of curcumin in *CL* have been demonstrated in multiple chronic inflammatory diseases such as, psoriasis, solar radiation and skin aging (Panahi et al., 2019) and yet the therapeutic efficacy of the *Curcuma Longa* essential oil (*CL*EO) especially on AD is not known.

Essential oils (EOs) treat dermatological conditions (Rajab, 2022). They are natural, do not damage the skin and increase skin penetration (Caliskan and Karakus, 2020).Their compounds have anti-inflammatory, and anti-itch properties (Vaughn et al., 2018). *Curcuma longa* essential oil(*CL*EO) is anti-inflammatory (Jaiswal and Naik, 2021) and repairs skin damage(Kashyap et al., 2022).

In this experimental study, the aim was to evaluate the anti-inflammatory effects of *Curcuma Longa* essential oil on Acetone induced Atopic dermatitis using albino wistar rats.

## Materials and Methods

### Collection and authentication of plant material

A Mature (10 months) whole plant was collected from an experienced Agriculturist in Western Uganda, Bushenyi district; Kyeizooba sub county; Rwanyankara village. It was dried in a herbarium for four weeks after collection; presented for identification and authenticated by a taxonomist Dr Olet Eunice; Department of Biology, Faculty of Science, Mbarara University of Science and Technology in Western Uganda with Voucher Number NN-2024-001.

### Extraction of *Curcuma longa* Essential oil (*CL*EO)

Fresh rhizomes (Ibáñez and Blázquez, 2021) of *Curcuma longa* were washed using running water, dried with paper towel and grated. 500g of grated *Curcuma longa* were loaded in a round bottomed flask in 1L of water and hydro-distilled using the Clevenger apparatus for 5 hours (Mc Gaw and Skeene, 2021). The *CL*EO was by extracted as the distillate. 40g of anhydrous sodium sulphate was added to the collection bottle (20ml) and then the collected pure essential oil added to fill the bottle. The sulphate removes the water remaining after the extractions (Avanço et al., 2017). The pure *Curcuma Longa* essential oil was then skimmed off the floral water from the bottle sing a dropper. The oil was stored at 4 - 8 °C and protected from light prior to us use (Avanço et al., 2017).

### Preparation of *CL*EO for experimental use

2ml of the *CL*EO was mixed with 98 ml of Olive oil prior to use as treatment for safety measures (Bensouilah Janetta, 2006).

### Laboratory Animals

They were kept in transparent glass cages measuring 12cm by 10cm in the well-ventilated animal house. They were acclimatized for seven days under good laboratory conditions (12 hours light/dark cycle; room temperature) before the start of the experiment. The animals were allowed to free access to standard rodent chow and water *ad libitum*.

### Induction of Atopic dermatitis

Animals were sedated with Di ethyl ether for 2 minutes. The dorsal part of the skin (4cm x 4cm) close to the tail was marked for use in the experiment and shaved using Veet cream. The shaving cream was then removed with a cotton ball containing saline water. The skin was then subjected to acetone using a soaked cotton wool for 5 minutes for three days (Barcelos et al., 2013). All the induced experimental animals were observed for itching and presence of red patches on the skin.

### Grouping of animals

The study used eight weeks old (Choi et al., 2020) old, 30 male and female albino Wistar rats; n=6 sourced from the animal facility of Kampala International University (KIU), Western Uganda. Group 1(Negative control) were not induced and not treated; Group 2 (Positive control) were induced and not treated; Group 3 were induced and the treated with 0.1% topical hydrocortisone; Group 4 were induced and treated with topical *CL*EO and Group 5 were induced and treated with both topical 0.1% Hydrocortisone and topical *CL*EO.

### Dosage

2% *CL*EO; Three drops were applied to the affected area and massaged for a minute. Hydrocortisone; A fingertip unit was applied to the affected area and massaged for a minute.

### Laboratory Analysis of AD

After 28 days of the experiment, the rats were starved overnight in preparation for dissection. Rats were euthanized with Di ethyl ether for 5 minutes. Small biopsies cut at 5 µm thickness were obtained from the dorsal skin of the rats where induction and treatment took place. Experimental rats were then humanely sacrificed and blood collected by cardiac puncture.

Technicians were blinded to the groups.

### Anti-pruritic effect of the treatment

The rats were kept in a transparent cage to allow use of a camera and asses scratching which is the result of itching. Observation was done in the morning for a 5 minutes and at night for 5 minutes and the results used for grading. The difference in the observation hours was aimed at creating a difference in the observation during active hours(night) and dormant hours(morning).The degree of scratching was quantified as the total number of bouts of scratching in the observation period (Lariosa-Willingham et al., 2023). This measured itching. A total score was taken on a weekly basis at the start of the week.

### Histological analysis; Mast cells

Histological examination of the skin tissues was carried out using standard H&E procedures. The skin Biopsies were received in neutral buffered formalin in a ratio of 1: 10 (tissue: formalin)They were processed, stained and analysed under a microscope together with a pathologist to give results indicating the differences in infiltration.

### Biochemical analysis; C - reactive protein

Samples were received in red top vacutainers and analysed using CRP ELISA kit.

### Quality control

Plant was transported in a cool box after collection. Rhizomes were kept in a refrigerator until extraction. The Rats were treated uniformly. Rats were fed with distilled water to minimise causes of infections. The skin biopsies were kept in neutral buffered formalin until histopathological analysis to maintain the tissue composition. A qualified experienced Pathologist guided the microscopic work. Manufacturer’s instructions were followed for the CRP Elisa kit.

### Statistical Analysis

Collected raw data organized in tables using Microsoft excel. Data were analyzed using Prism software (Graph-Pad Software; San Diego, CA).Pruritis and CRP results were expressed as mean ± SEM. Multiple group differences were analysed at 95% confidence interval through the one-way Analysis of Variance (ANOVA). Differences between two groups was by student’s t test.

## Results

### Anti pruritic effect of *Curcuma Longa* Topical Essential oil against acetone induced Atopic Dermatitis

The frequency of scratching was observed for 10 minutes on a weekly basis. It gradually reduced until it stopped. Scracthing measured pruritus in this experiment.

**Figure 1.**
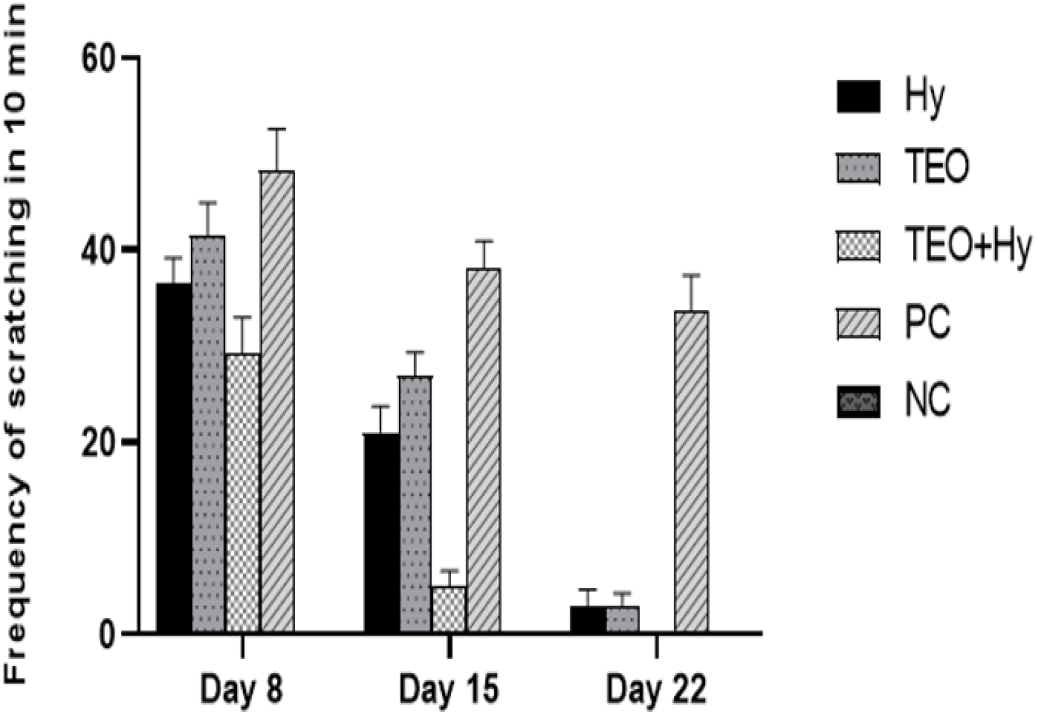
Anti pruritic effect of the treatments on acetone induced AD. Hy-indicates the group treated with 0.1% hydrocortisone ointment (HYCORT); TEO – indicates the group treated with *CL*EO; TEO+ Hy – indicates the group treated with both Hydrocortisone and *CL*EO; PC – indicates Positive control; Nc – indicates Negative control. Data are shown as mean ± S.EM (n = 6)

**Figure 2.**
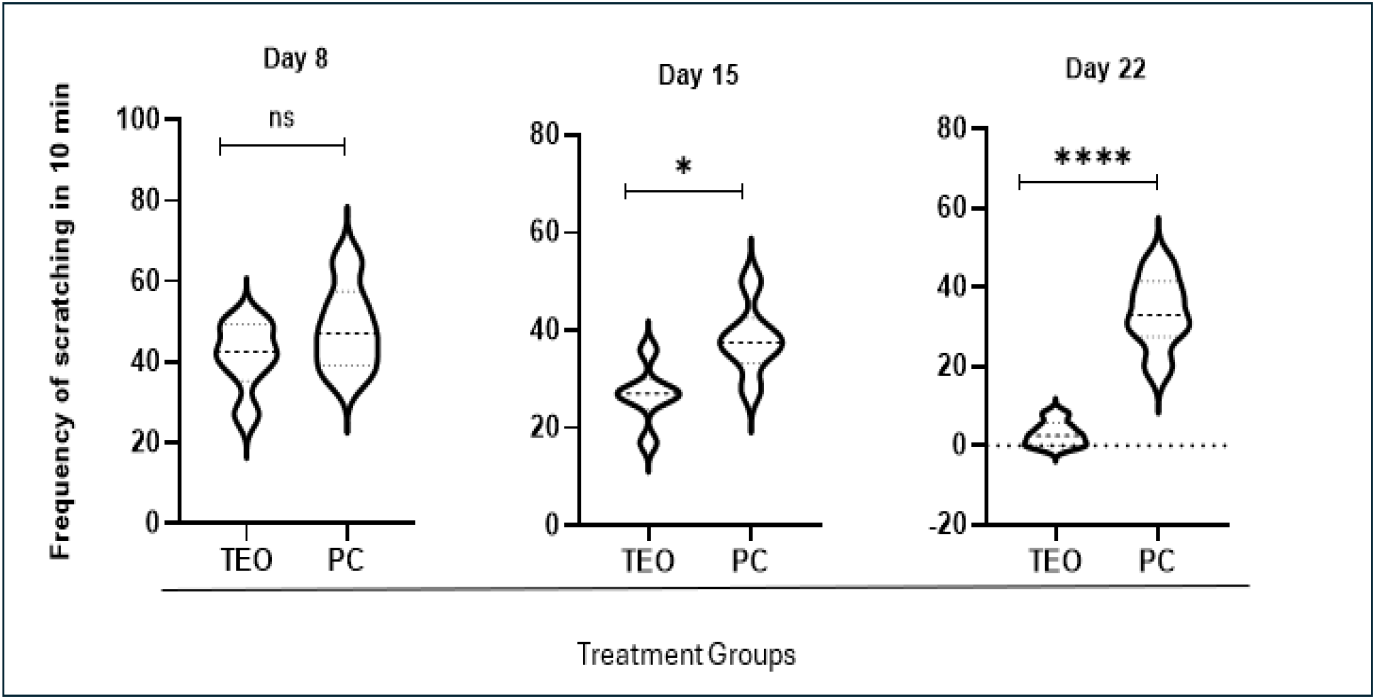
Comparison of the pruritus in the two groups: The TEO (Group treated with CLEO) and the positive control group on the analysis days. On day 8, there was no significant difference (P = 0.242). On day 15, there was a small significant difference (P<0.05; *). On day 22, there was an increased significant difference (P<0.0001) (****)

### Histo-pathological analysis; Mast cells

**Table 1.**
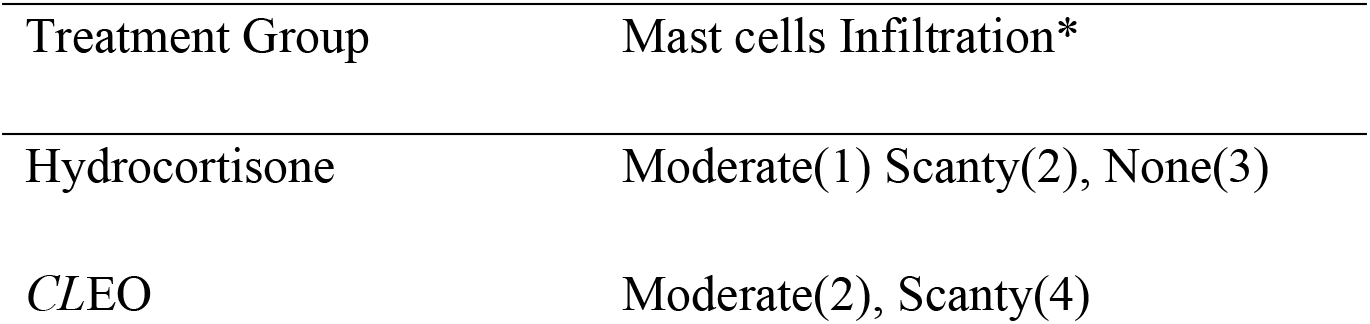
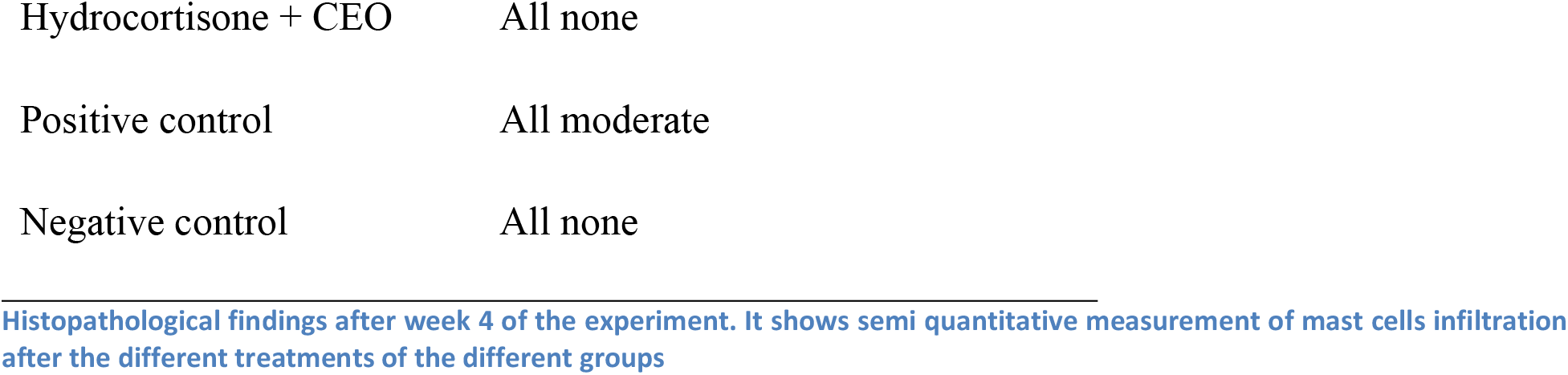
Infiltration of Mast cells.

| Treatment Group | Mast cells Infiltration* |
| --- | --- |
| Hydrocortisone | Moderate(1) Scanty(2), None(3) |
| CLEO | Moderate(2), Scanty(4) |
| Hydrocortisone + CEO | All none |
| Positive control | All moderate |
| Negative control | All none |

Histopathological findings after week 4 of the experiment. It shows semi quantitative measurement of mast cells infiltration after the different treatments of the different groups

**Figure 3.**
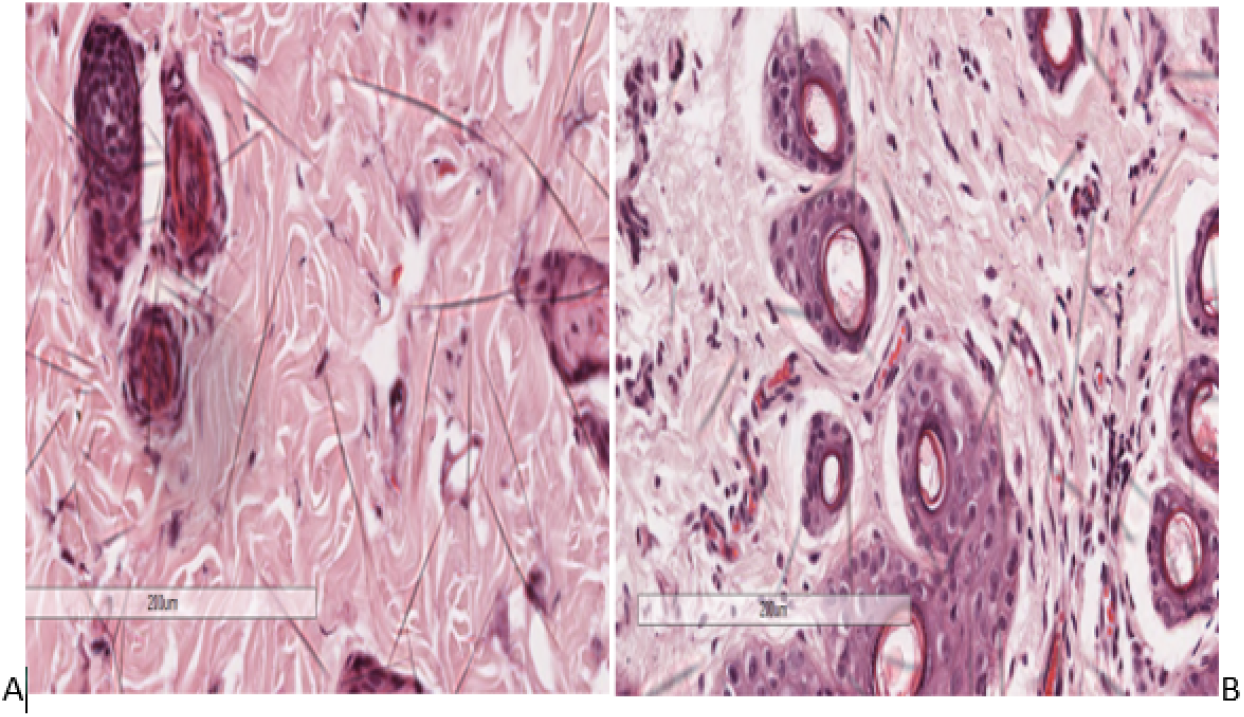
Hematoxylin and Eosin stain on histological sections of skin tissue showing infiltration of stained mast cells as observed under a microscope at magnification 200µm ; A (group treated with *CL*EO) Infiltration of mast cells is scanty. B (Positive control-group induced but not treated) Infiltration is Moderate.

### Biochemical analysis

The CRP Levels were significantly different when descriptive statistics were ran using graph pad prism; P <0.0001; f= 81.30. *CL*EO is also referred to as TEO. Dunnet’s multiple comparison tests with in groups demonstrated a significant difference between Positive Control and Negative Control with a mean difference of -0.02633. Treatment group of *CL*EO and Positive group were significantly different with p value 0.0001.

Key; TEO; *Curcuma Longa* Essential oil, Hy-Hydrocortisone; No Rx-Positive control (induced but not treated) NC – Negative Control (not induced and not treated)

**Figure 4.**
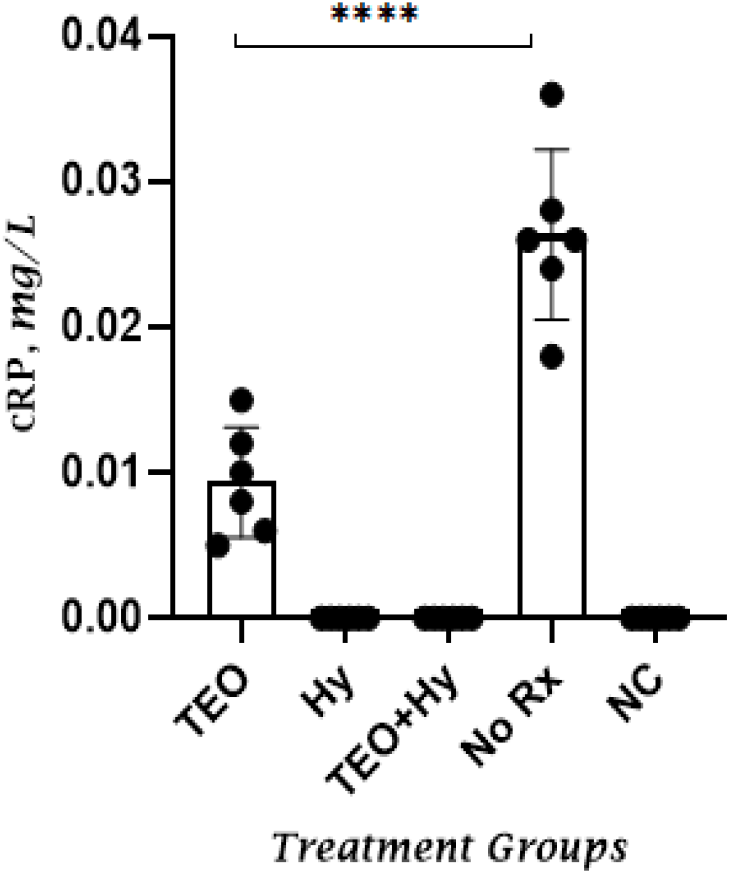
A graph showing the difference in means of CRP values in the different treatment groups ; TEO-CLEO treatment; HY-Hydrocortisone treatment; No Rx-Negative Control; not treated and induced NC-Normal Control; not treated and not induced.

**Figure 5.**
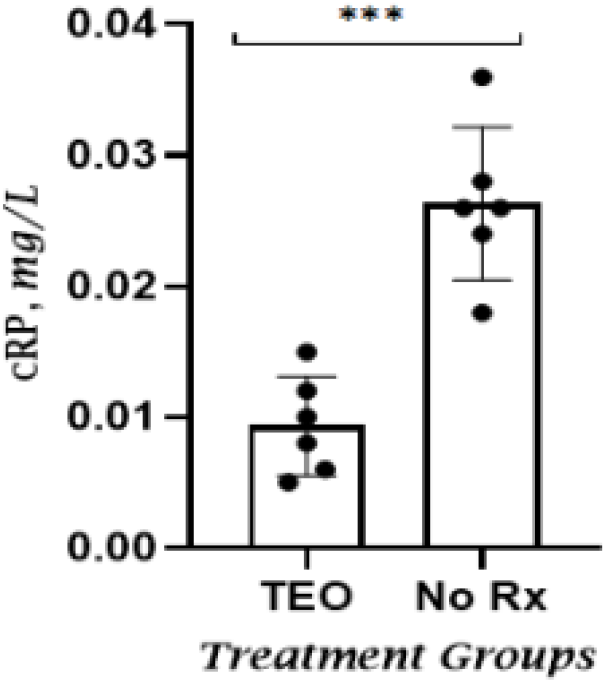
Graph showing the difference in means of CRP values between the the group that received the Topical *CL*EO and the negative control. Crp levels are significantly different in the group treated wth *CL*EO (TEO)which indicated the anti inflammatory effects of the essential oil. TEO-Group treated with CLEO; Negative control-Group induced and not treated.

## Discussion of results

### *CL*EO and pruritis

The pruritis greatly reduced and eventually stopped in the treatment group that had *CL*EO. The Scratching frequency was used to measure itching. There was a significant decrease in the scratching frequency in the group treated with CLEO compared to the positive control with P <0.0001 (^****^). Similarly peppermint essential oil treats pruritus in pregnant women. Essential oils exhibit anti pruritic activity by suppressing the expression NFκB (Mohajerani et al., 2019). NF-κB is the main transcriptional factor that regulates the expression of pro-inflammatory mediators. Terpene-induced reduction of NF-κB activity is the direct cause of inhibition of expression of pro-inflammatory mediators (Kim et al., 2020). *CL*EO consists of terpenes (Kim et al., 2020) and therefore may down regulate NF-κB activity. Similarly, topical *Mentha arvensis* essential oil showed anti-inflammatory effects in a dinitrochlorobenzene (DNCB)-induced murine model of AD by inhibition of the NF-κB (Kim et al., 2021).The application of terpenes and terpenoids to alleviate inflammation has been shown successful in the mitigation of AD (Masyita et al., 2022). Essential oils exhibit anti pruritic activity by stabilizing mast cells (Mohajerani et al., 2019). *CL*EO stabilizes Mast cells (de Sousa et al., 2023). Terpenes constitute the composition of the *CL*EO and therefore It may treat pruritus in AD.

### *CL*EO and Mast cells

*CL*EO decreased the Mast cell infiltration in the skin biopsies.*CL*EO inhibits Itching/ pruritis as explained above. This in turn inhibits Mast cell degranulation. Mast cell cell degranulation is an inflammatory response as a result of the itching. Increase in Mast cell degranulation results to an increased infiltration observed under a microscope after H & E staining. *CL*EO stabilizes Mast cells (de Sousa et al., 2023); therefore may stop inflammation in AD. Similarly, Geranium Essential oil showed anti-inflammatory effects by inhibiting the degranulation of cultured mouse mast cells (Kobayashi et al., 2016).

### *CL*EO and CRP levels

*CL*EO reduced CRP levels. The CRP levels in the group treated with the *CL*EO were significantly lower than the levels in the group that was induced and not treated with a P value 0.0001. Mast cells activated in AD release a large number of inflammatory cytokines such as IL-6 and TN-alpha through degranulation (Siiskonen and Harvima, 2019). Through the factors produced on activation, MCs regulate the recruitment, transport, and functions of cells taking part in the skin immune response in AD. IL-6 in circulation stimulates production of CRP in the liver and release into the blood stream (Del Giudice and Gangestad, 2018). *CL*EO constitutes terpenes which promote mast cell stabilization (de Sousa et al., 2023) and in turn reduce the release of CRP.

### *CL*EO and dry skin

AD is a dermatological condition with a dehydrated skin due to increased TEWL (Douladiris et al., 2023). *CL*EO is a very good moisturizer (Barbalho et al., 2021). Moisturizers regulate of SC water content with their different hydration capacities and have been used to treat AD (Douladiris et al., 2023). *CL*EO repairs skin damage (Kashyap et al., 2022) yet the most instrumental problem in AD is skin barrier function due to damage. Atopic dermatitis causes an itching/ pruritis response which is neurogenic, scratching and thereby mechanical damage of the skin barrier (Martel et al., 2017). Restoration of the skin’s barrier, focused on the restoration of the SC, is the essence of management of AD (Douladiris et al., 2023). Moisturizers maintain the integrity of the epidermal barrier and promote its protective function against dehydration (Elmariah and Lerner, 2011). *CL*EO being a moisturizer may therefore help to treat dry skin. During the study, the group that was treated with the CLEO had smooth moisturized skin compared to the Negative control. This was an observation taken on the day of sacrificing.

## Conclusion and Recommendation

Our results suggest that *Curcuma longa* Topical essential oil has therapeutic potential on treat Atopic dermatitis. Further studies should be considered for phytochemical analysis *of CLEO* from western region in Uganda and 2% *CL*EO should be evaluated in a clinical trial.

## Acknowledgements

We acknowledge contributions of laboratory technician at Mbarara University of science and technology and Kampala International University including Mr. Mutekanga Emmanuel and Mr Charles

## Authors’ contributions

DT, PMA and NN developed the topic, designed the experiment, NN carried out laboratory work and biochemical analysis under guidance of a technician, AB carried out histopathological analysis and KM analysed data. NN developed the first Manuscript, the whole group revised and contributed to the final manuscript.

## Data Availability

All data generated or analyzed during this study is to be included in the published article.

## Ethics

This study was approved by Mbarara University of Science and Technology Faculty of Medicine Research Committee and the Institutional Animal Care and Use Committee (IACUC) Makerere University, Uganda with reference number SVAR_IACUC/166/2023. The animals were treated according to the guidelines of Uganda National Council of Science and Technology (UNCST) 2021 such as freedom of expressing behavior, consent from the owner about what the animals are going to be used and the use of the 3R’s to use minimum number of rats possible.

